# F-type Pyocins are Diverse Non-Contractile Phage Tail-Like Weapons for Killing *Pseudomonas aeruginosa*

**DOI:** 10.1101/2021.02.16.431561

**Authors:** Senjuti Saha, Chidozie D. Ojobor, Erik Mackinnon, Olesia I. North, Joseph Bondy-Denomy, Joseph S Lam, Alexander W. Ensminger, Karen L. Maxwell, Alan R. Davidson

**Author notes:** Address correspondence to Alan R. Davidson.

## Abstract

Most *Pseudomonas aeruginosa* strains produce bacteriocins derived from contractile or non-contractile phage tails known as R-type and F-type pyocins, respectively. These bacteriocins possess strain-specific bactericidal activity against *P. aeruginosa* and likely increase evolutionary fitness through intraspecies competition. R-type pyocins have been studied extensively and show promise as alternatives to antibiotics. Although they have similar therapeutic potential, experimental studies on F-type pyocins are limited. Here, we provide a bioinformatic and experimental investigation of F-type pyocins. We introduce a systematic naming scheme for genes found in R- and F-type pyocin operons and identify 15 genes invariably found in strains producing F-type pyocins. Five proteins encoded at the 3’-end of the F-type pyocin cluster are divergent in sequence, and likely determine bactericidal specificity. We use sequence similarities among these proteins to define 11 distinct F-type pyocin groups, five of which had not been previously described. The five genes encoding the variable proteins associate in two modules that have clearly re-assorted independently during the evolution of these operons. These proteins are considerably more diverse than the specificity-determining tail fibers of R-type pyocins, suggesting that F-type pyocins emerged earlier or have been subject to distinct evolutionary pressures. Experimental studies on six F-type pyocin groups show that each displays a distinct spectrum of bactericidal activity. This activity is strongly influenced by the lipopolysaccharide O-antigen type, but other factors also play a role. F-type pyocins appear to kill as efficiently as R-type pyocins. These studies set the stage for the development of F-type pyocins as anti-bacterial therapeutics.

**IMPORTANCE:** *Pseudomonas aeruginosa* is an opportunistic pathogen that causes a broad spectrum of antibiotic resistant infections with high mortality rates, particularly in immunocompromised individuals and cystic fibrosis patients. Due to the increasing frequency of multidrug-resistant *P. aeruginosa* infections, there is great interest in the development of alternative therapeutics. One alternative is protein-based antimicrobials called bacteriocins, which are produced by one strain of bacteria to kill other strains. In this study, we investigate F-type pyocins, bacteriocins naturally produced by *P. aeruginosa* that resemble non-contractile phage tails. We show that they are potent killers of *P. aeruginosa*, and distinct pyocin groups display different killing specificities. We have identified the probable specificity determinants of F-type pyocins, which opens up the potential to engineer them to precisely target strains of pathogenic bacteria. The resemblance of F-type pyocins to well characterized phage tails will greatly facilitate their development into effective antibacterials.

## INTRODUCTION

With increasing antibiotic resistance, there is a strong incentive to identify alternative anti-bacterial therapeutics. To this end, interest in using phages or parts of phages to treat bacterial infections has greatly increased in recent years (1), and phage treatments have proven effective in clearing bacterial infections in humans (2-4). This success notwithstanding, there are potential drawbacks to phage therapy, including the possibility that introduced phages may acquire and transmit virulence or antibiotic resistance genes, and that negative outcomes may arise from long-term phage reproduction within a patient. To circumvent these problems, the therapeutic potential of phage tail-like bacteriocins, also referred to as tailocins, is also being explored. Tailocin encoding operons, which are found in many diverse bacterial species, are likely derived from prophages. The utility of tailocins as antibacterials has been amply demonstrated (5, 6). Like phages, tailocins are highly specific for their target organism, but they possess additional advantages. A single tailocin type can be engineered to kill a variety of bacterial species (7, 8), and tailocins can be efficiently produced in easily cultured organisms, such as *E. coli* (9) or *B. subtilis* (10). In this work, we provide a detailed investigation of a group of tailocins produced by *Pseudomonas aeruginosa* that are related to non-contractile phage tails.

The tailocins of *P. aeruginosa*, discovered many decades ago (11), fall into two types known as F-type pyocins and R-type pyocins. All *P. aeruginosa* strains possess a gene cluster located between the *trpE* and *trpG* genes encoding F-type, R-type or both types of pyocins. R-type pyocins, which are related to contractile-tailed phages, such as *E. coli* phage P2 (12), have been extensively studied. These entities are produced by different strains of *P. aeruginosa* and have the ability to kill other strains of the same species. R-type pyocins bind specifically to target strains, and then puncture their inner membrane, leading to rapid cell death (13). Derivatives of R-type pyocins with engineered tail fibres are able to kill other species of bacteria, such as *E*.*coli* and *Yersinia pestis*, and these engineered variants have shown efficacy in preventing and/or ameliorating infection in animal models (6, 9, 14, 15). F-type pyocins, which are related to non-contractile tailed phages, such as *E. coli* phage lambda (12), have been studied much less than the R-type. Although encoded in more than half of *P. aeruginosa* strains (16), no experimental work has been published on F-type pyocins since 1981 (17). The activities of F-type pyocins produced by five different strains have been described in the literature (17-20), each of which killed distinct sets of *P. aeruginosa* strains. However, the sequences of the operons encoding only two of these are known. Based on genome sequencing data, four further groups have been defined (16), but neither the production nor activity of these groups was assessed. The mechanism of action and killing specificity determinants of F-type pyocins have not been defined.

Given the potential importance of tailocins in treating bacterial infections and the relative dearth of information pertaining to F-type pyocins, we undertook a comprehensive investigation of F-type pyocins encoded in a large number of *P. aeruginosa* strains. The goals of this study were to bioinformatically characterize F-type pyocin operons and correlate sequence diversity with the killing spectra of defined F-type pyocin groups. Through this process, we have identified 11 distinct groups of F-type pyocins, and their likely specificity determinants. We conclude that these F-type pyocins have the potential to be engineered as highly effective anti-bacterial therapeutics.

## RESULTS

### Selection of *P. aeruginosa* strains for this study

To gain an understanding of the diversity of R-type and F-type pyocins produced by *P. aeruginosa*, we produced lysates of diverse strains selected from our collection (21) by treating cultures with mitomycin C, which induces pyocin production and cell lysis (22). Each of these lysates was examined by transmission electron microscopy (TEM), and those displaying abundant levels of R- and/or F-type pyocins were further analyzed (Fig. 1). Ultimately, a set of 30 strains was chosen that produced only F-type (n = 8), only R-type (n = 9) or both R- and F-type (n = 13) pyocins. This set contained clinical and environmental strains from seven different countries, collected over a few decades (Supplementary Table 1). Twenty-eight of the 30 strains were sequenced, assembled and annotated in this study (annotated genomes of strains PAO1 and PA14 were obtained from the Pseudomonas Genome Database (23)).

**FIG 1.**
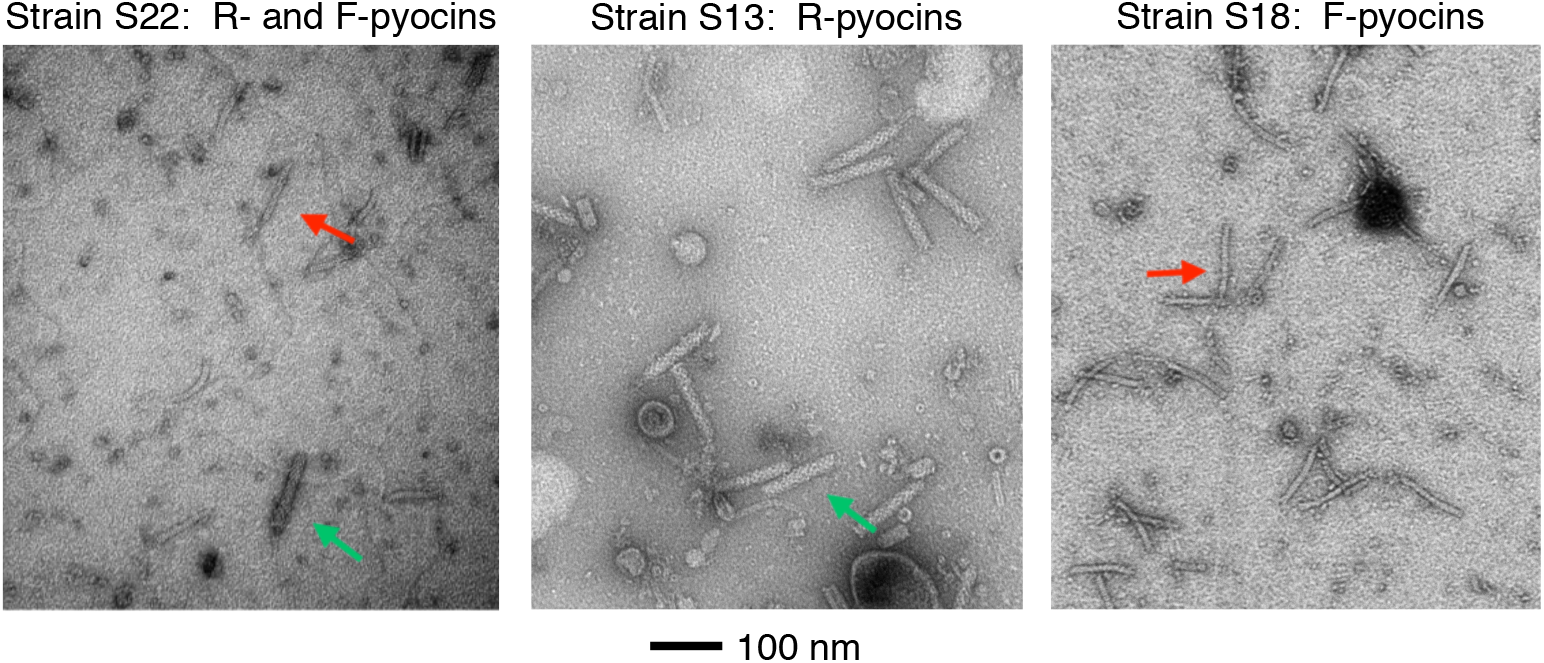
Transmission electron micrographs of lysates of cells producing R- and F-type pyocins. Shown are a lysate of strain S22 (left panel), which produces both R- and F-type pyocins; a lysate of strain S13 (middle panel), which produces just R-pyocins; and a lysate of strain S18 (right panel), which produces just F-pyocins. R-type pyocin particles are indicated by green arrows and F-type pyocin particles are indicated by red arrows. Grids were negatively stained with uranyl acetate. The scale bar shown applies to all three micrographs.

### Conserved features of R/F-type pyocin clusters

The R- and F-type pyocin gene clusters are invariably found between the *trpE* and *trpG* genes in *P. aeruginosa* (12). To locate these gene clusters in each genome that was sequenced in this study, the regions between gene *trpE* and *trpG* were extracted and analyzed (Fig 2a). Each of the 30 sequenced genomes was found to encode an F-type- or R-type pyocin, or both, corresponding with the observed production of pyocin particles observed by electron microscopy. In total, 23 R-type and 21 F-type pyocin gene clusters were present in our analyses. The gene content of the pyocin clusters was constant across all the strains. We observed eight genes, designated *pyoRF1* to *pyoRF8*, which were found in all clusters, 15 genes specific to R-type pyocins (*pyoR1* to *pyoR15*), and 15 genes specific to F-type pyocins (*pyoF1* to *pyoF15*) (Fig. 2a, Table 1). The *pyoRF1* and *pyoRF2* genes encode the PrtN activator and PrtR repressor, respectively. These proteins regulate expression of the cluster in response to DNA damage as previously described (22). The *pyoRF3* and *pyoRF4* gene products are uncharacterized, but their predicted functions suggest a role in regulating expression of the gene cluster. Analysis by HHpred indicates that the *pyoRF3* gene encodes a putative zinc-binding transcription factor and *pyoRF4* encodes a putative transcription anti-terminator protein similar to gpQ of phage lambda (24). Homologs of PyoRF3 are found in more than 100 phage and prophage genomes, while homologs of PyoRF4 are found in a much smaller number of phages and prophages. The pyoRF5 to pyoRF8 genes encode a complete set of phage-like lysis genes, including a peptidoglycan hydrolase, holin, and Rz and Rz1-like spannins (25). In operons encoding only F-type pyocins all eight *pyoRF* genes precede the genes encoding the F-type pyocin specific genes. In clusters encoding just R-type pyocins, or those encoding both R- and F-type pyocins, the R-type pyocin specific genes are inserted within the lysis gene cluster between *pyoRF5* and *pyoRF6* (Fig. 2a).

**FIG 2.**
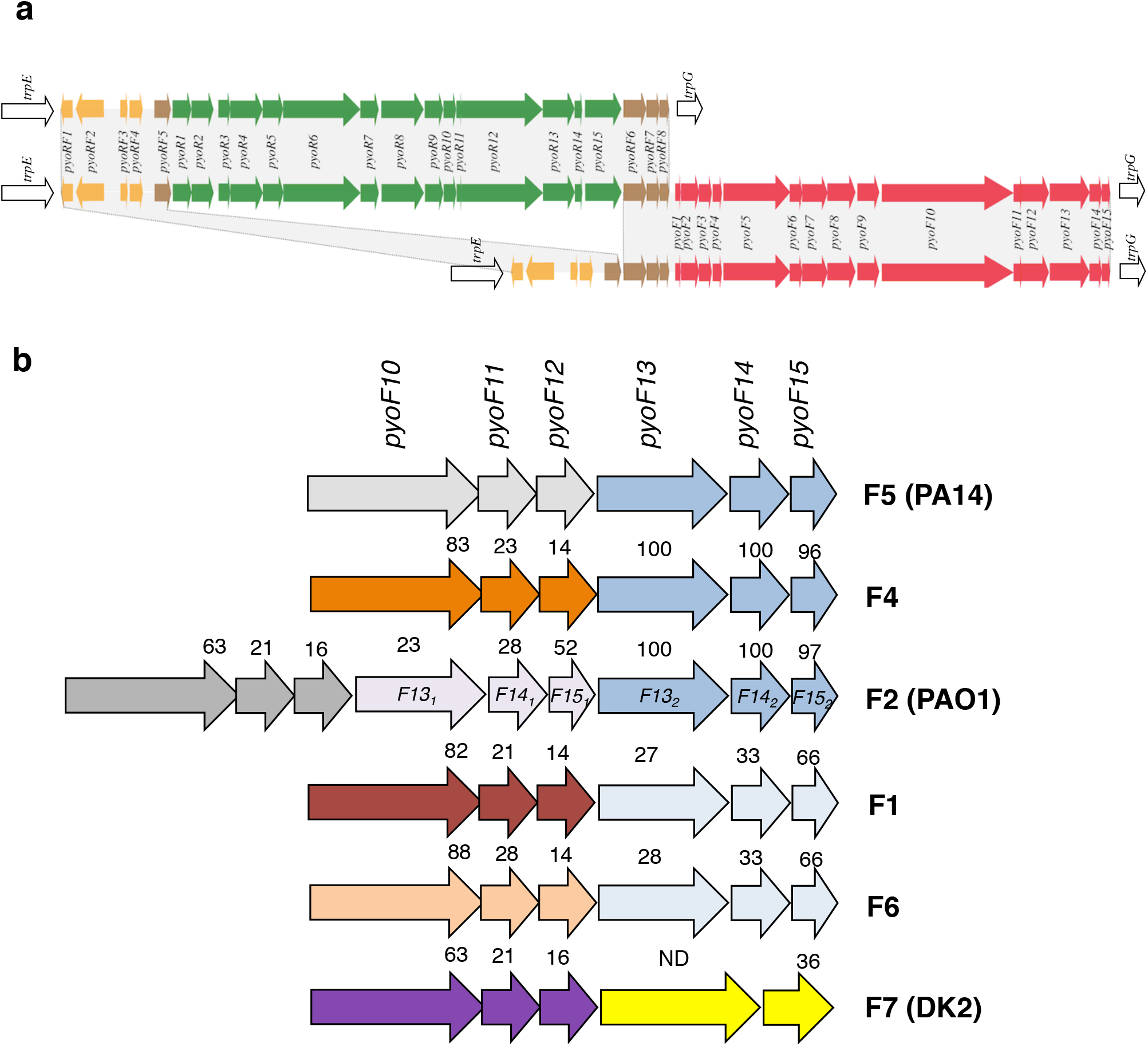
R- and F-type pyocin operons. (a) Three types of R- or F-pyocin operons are found in *P. aeruginosa*: operons encode just R-pyocins (top), R- and F-pyocins (middle, or just F-pyocins (bottom). All three types share the same regulatory genes (orange) and lysis genes (brown). Genes unique to R-pyocins are colored green and those unique to F-pyocins are colored red. All three types of operons are located in the same position in the *P. aeruginosa* genome between the *trpE* and *trpG* genes. (b) A close up of genes encoded at the 3’-end of the F-type pyocin clusters show the six different groups identified in our sequenced strains. The numbers above the genes indicate the percent pairwise sequence identity of the encoded protein with the homolog found in strain PA14 (group 5). Proteins PyoF13, PyoF14 and PyoF15, which are duplicated in group F2, are very closely related (>95% sequence identity) in groups F5, F4 and F2 (second group) as indicated by their coloring. The same proteins are highly similar in groups F1 and F6. In the cases of PyoF10 and PyoF13, sequence comparison were performed including only their variable C-terminal domains. *P. aeruginosa* strains where certain groups were previously identified are shown in parentheses.

**Table 1.**
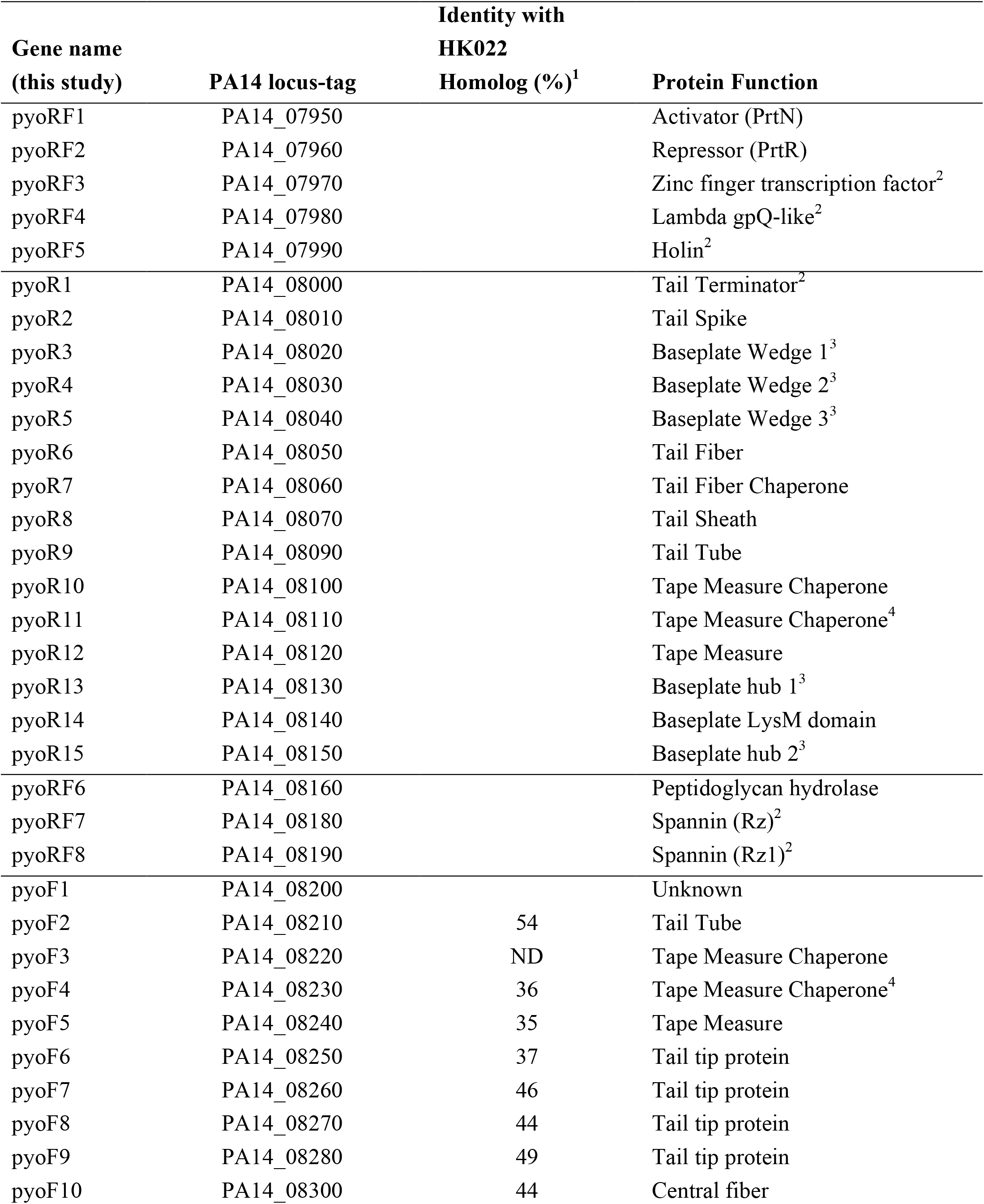
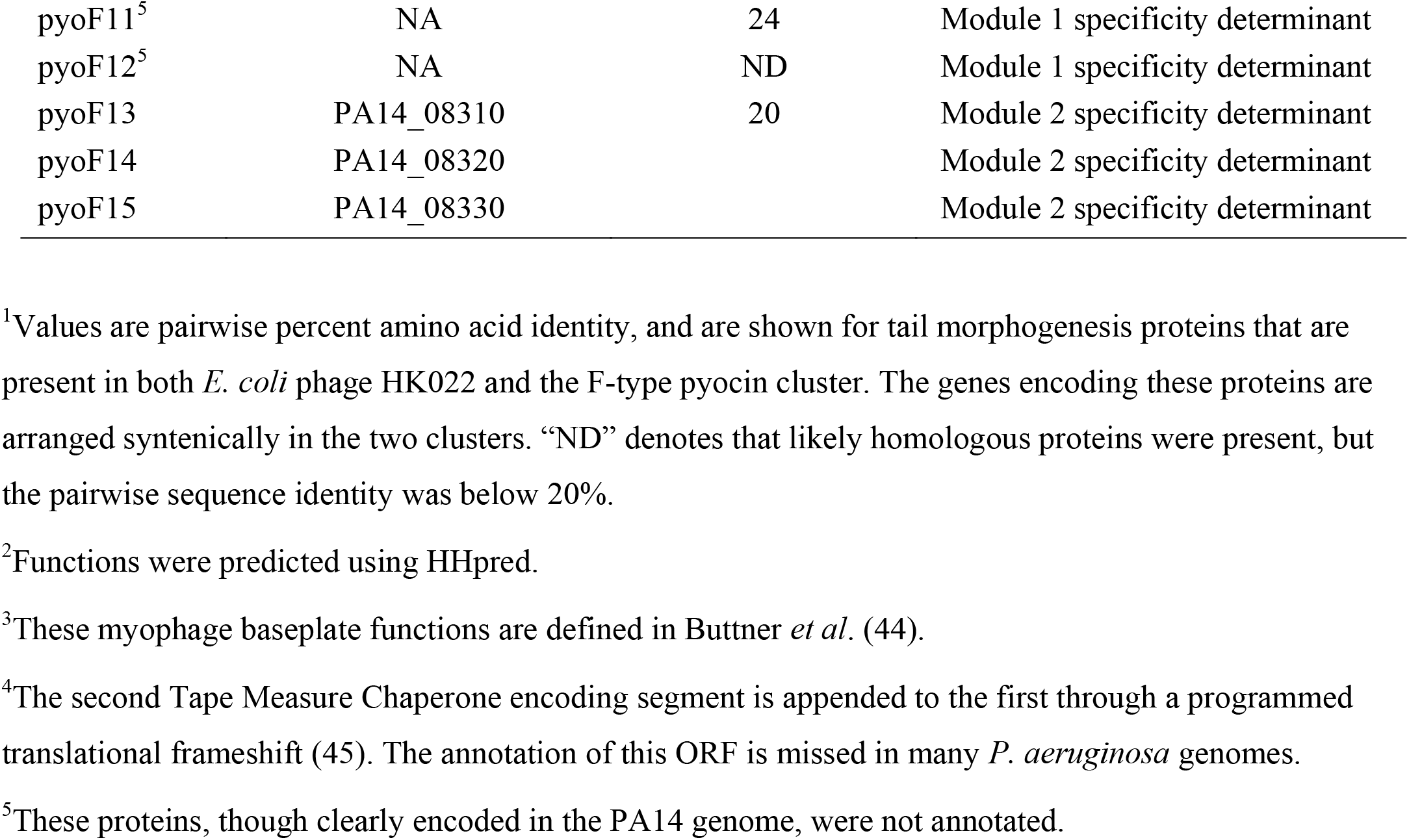
Gene composition of the pyocin operon in *P. aeruginosa* strain PA14.

Within the 22 R-type pyocin clusters in the genomes studied here, 13 of the 15 encoded proteins are highly conserved among the clusters, with at least 97% sequence identity between each gene product, as has been previously documented (15). The two proteins that vary significantly are PyoR6 and PyoR7, which encode the tail fiber and tail fiber chaperone, respectively. The tail fiber determines the specificity of R-type pyocins and the chaperone is specific to its cognate fiber. We compared the fiber and chaperone proteins of each of our sequenced clusters to those of the characterized R-type pyocin types (15). Fiber sequences of the R2-, R3- and R4-types are very similar to each other (> 98% identical). Hence, we considered groups R2, R3 and R4 as one group and called it group R2, as was done in a previous study (26). For the R-type pyocin clusters sequenced here, ten belonged to the R1 group, ten to the R2 group, and two to the R5 group. As R-type pyocins have been well characterized in previous studies (7, 8, 15), we focused the present investigation on the F-type pyocins.

### Conserved proteins encoded in the F-type pyocin cluster

The *pyoF2* to *pyoF10* genes encode confidently annotated functions required for formation of the F-type pyocin tube and tip (Table 1). The protein products of each of these genes are clearly homologous to phage tail proteins (27), and these proteins are very similar (∼ 95% pairwise sequence identity) among the 21 F-type pyocin gene clusters that we have analyzed. Although the F-type pyocin genes are arranged in an order that is syntenic with the genome of the well characterized *E. coli* phage lambda (12), only the tail tip and central fiber proteins (PyoF6 to PyoF10) of this phage share significant sequence identity with F-type pyocin proteins (31 to 38% sequence identity). The phage tail region with the greatest similarity to the F-type pyocin cluster across the tube and tail tip region is that from *E. coli* phage HK022. (Table 1). The HK022 proteins share 43% sequence identity, on average, to those of the F-type pyocin (Table 1). No prophage tail region was more closely related to the F-type pyocin cluster than phage HK022 across the whole cluster, though some *P. aeruginosa* prophages were more closely related to the 3’-end of the cluster where genes encoding the tail tip proteins are located.

An unusual feature of F-type pyocin regions as compared to phage tails is the lack of any protein with detectable similarity to a tail terminator. This protein is essential for phage tails because it is required to join the tail to the head (28). The tail terminator also prevents uncontrolled polymerization of the tails of some (28), but not all phages (29). Since F-type pyocins are not joined to a head, the tail terminator appears to be dispensable. The *pyoF1* gene lies in the genomic position expected for a tail terminator gene. However, the 95 amino acid protein encoded by this gene bears no detectable sequence similarity to tail terminators, has no homologs outside of *P. aeruginosa* F-type pyocin clusters, and stop codons are observed in this ORF in several strains. Thus, we conclude that this is not a functioning protein as was also concluded in a previous publication (12).

### F-type pyocins can be grouped based on proteins encoded at the 3’-end of the cluster

The host range specificity of phages is determined by proteins located at the tail tip, which are typically encoded by genes at the 3’-end of tail-encoding regions (27). The analogous proteins in the F-type pyocin are encoded by genes *pyoF10*-*pyoF15*. Non-contractile tails resembling F-type pyocins possess a long (> 700 residues) central fibre protein that projects directly below the tail tip. In phage lambda, the region within the last 250 residues of the central fiber mediates host cell specificity and surface binding (30, 31). The homologous protein in F-type pyocins is encoded by *pyoF10*. We observed that the first 1160 residues of the PyoF10 proteins are highly conserved among F-type pyocins (> 93% sequence identity), but the last 60 residues vary greatly, with pairwise identities in this region often ranging below 35% (Fig. S1). This sequence variability is consistent with a role for the C-terminus of PyoF10 in mediating host specificity.

In addition to the last 60 residues of PyoF10, the other five proteins encoded at the 3’-end of the F-type pyocin cluster, PyoF11 to PyoF15, were found to vary considerably in sequence between different F-type pyocin clusters. Based on pairwise comparison of homologous proteins encoded in this region of the clusters (Fig. S1), the F-type pyocin regions found in different genomes were divided into six groups, F1, F2, F4, F5, F6 and F7 (Fig. 2b). Regions were placed into the same group if each of their corresponding homologous proteins shared at least 90% sequence identity with all others in the group. The nomenclature used here extends from previous work where groups F1 to F3 were established based on differences in host killing specificity (17). We do not know if any of the groups identified here match group F3 because no examples from this group have been sequenced. The groups that we called F4 and F6 have not been previously recognized, while group F5 and F7 were previously described in *P. aeruginosa* strains PA14 and DK2, respectively (16). The two most frequently occurring groups are F2 (11 members) and F7 (4 members). The F1, F5 and F6 groups were encoded only in pyocin clusters that also encoded R-type pyocins, while F4, F7, and F2 group clusters were found in the absence of R-type clusters except in two instances (both F2 group). Further bioinformatic comparisons described below compare representative protein sequences from each of the six F-type pyocin groups that we identified here.

### PyoF11 and PyoF12 are newly recognized conserved proteins

PyoF11 and PyoF12 are proteins of unknown function that are encoded in every F-type pyocin region. These are the most diverse proteins in the F-type pyocin clusters, often displaying pairwise sequence identities of less than 25% (Fig. S1). Despite their diversity, the homologs of these proteins from the six groups could be convincingly aligned (Fig. S2a,b). We used HMMer (32) to create Hidden Markov Model (HMM) profiles from the PyoF11 and PyoF12 alignments. Searching with these HMMs, we identified more than 50 occurrences each of *pyoF11* and *pyoF12* gene homologs in diverse phages and prophages. These genes often occur together and invariably lie immediately 3’ to the central fiber gene (homolog of *pyoF10*). In some phage genomes, the *pyoF11* and *pyoF12* genes are very likely the last genes in the tail operon as they are followed immediately by lysis genes (e.g. *Burkholderia* phage Bcep176 and *Xanthomonas* phage CP1). These observations suggest that PyoF11 and PyoF12 function in conjunction with the central fiber protein, possibly binding to it or acting as chaperones to aid in folding of the fiber. PyoF11 and PyoF12 had not been previously recognized as conserved proteins in the F-type pyocin cluster because these ORFs are not annotated as proteins in most *P. aeruginosa* genomes. This is likely a result of the lack of annotation of these genes in the PA14 genome, which is commonly used as the reference genome for genome assembly and annotation. The functions of PyoF11 and PyoF12 homologs in phages have never been investigated.

### PyoF13, PyoF14, and PyoF15 are likely involved in host specificity

In addition to the central fiber, most non-contractile tailed phages possess genes downstream of the central fiber gene that also encode cell surface receptor binding proteins These are known as “side fibers” in *E. coli* phage lambda (33). The PyoF13 proteins, which share a common genomic position with the lambda side fibers, are likely involved in determining host range specificity, functioning as receptor binding proteins. A striking feature of the Pyo13 homologs is that their N-termini (the first 140 residues) are very similar among the F-type pyocin groups with pairwise sequence identities ranging from 55% to 90% while the pairwise identities in their C-terminal regions generally range between 20% and 35% (Fig. S1, S3). We surmise that the more conserved N-terminus of PyoF13 mediates binding of this putative receptor binding protein to the F-pyocin tail tip, while the variable C-terminus mediates cell surface binding. The fibers from different groups of R-type pyocins, which have been shown to determine bactericidal specificity (15), display the same type of conservation pattern with N-terminal regions (first 450 residues) displaying greater than 95% pairwise sequence identity and C-terminal regions (last 250 residues) displaying pairwise sequence identities between 50 and 70% (Fig. S4). In contrast to the F-type pyocins, there are only three distinct groups of R-type pyocins, as defined by fiber sequences, and there is much less variability.

Homologs of PyoF13, which share sequence similarity with its N-terminal region, are found in diverse phages and prophages and are located in similar genomic positions as *pyoF13*, downstream from the central fiber gene. Genes encoding homologs of PyoF14 (∼100 residues) and PyoF15 (∼75 residues) are also found in many phages and prophages, and they are invariably located downstream of *pyoF13* homologs or genes encoding other putative phage receptor-binding proteins. The sequences of PyoF14 and PyoF15 are variable, mirroring the sequence variation seen in the C-terminal regions of PyoF13 (Fig. S1). We expect that PyoF14 and PyoF15 are involved in host range specificity through interactions with PyoF13, or possibly by acting as chaperones for the assembly of PyoF13 as is required for phage-encoded receptor binding proteins (34).

Two F-type pyocin groups deviate from the others in the *pyoF13* to *pyoF15* region. The F2 group has a complete duplication of this region so that it possesses two copies of each gene. The proteins encoded by the first copy, PyoF13_1_ to PyoF15_1_, are distinct in sequence compared to the homologs in other groups (Fig. S1, Fig S3). Conversely, PyoF13_2_ to PyoF15_2_ are very similar (> 90% identical) to homologs found in groups F4 and F5. In contrast to all other groups, group F7 lacks a *pyoF14* gene. Consistent with this absence, its PyoF13 homolog displays a C-terminal region that has no detectable sequence similarity to the other groups.

### Characterization of F-type pyocin bactericidal specificity

To determine if our bioinformatic groupings of the F-type pyocin clusters correlate with bactericidal specificity, we examined the killing profiles of lysates produced from the 30 strains following induction by mitomycin C. Serial dilutions of lysates of each of the 30 strains were spotted onto lawns of the same 30 strains to produce an all-against-all matrix. Bactericidal activities were detected as zones of clearing on the bacterial lawn. Analysis of these data was complicated because *P. aeruginosa* produces other bactericidal entities in addition to R-type and F-type pyocins, including S-type pyocins (11) and bacteriophages. Since the presence of any of these can produce zones of clearing, further analyses were necessary to delineate the type of activity present. Testing serial dilutions of lysates allowed us to distinguish clearings produced by phages from those produced by pyocins (Fig. 3a). Due to their replicative nature, clearings resulting from phage lysates resolved into individual plaques upon dilution, while the clearings resulting from pyocins gradually disappeared without the appearance of individual plaques (Fig. 3a). Lysates were also spotted onto bacterial lawns containing proteinase K, which eliminated clearings caused by protease sensitive S-type pyocins (Fig. 3a) (11). By analyzing the activities of the 30 lysates on 30 strains in this manner, we detected more than 450 bactericidal combinations and found that greater than 90% were due to R- or F-type pyocins (Fig. S5).

**FIG 3.**
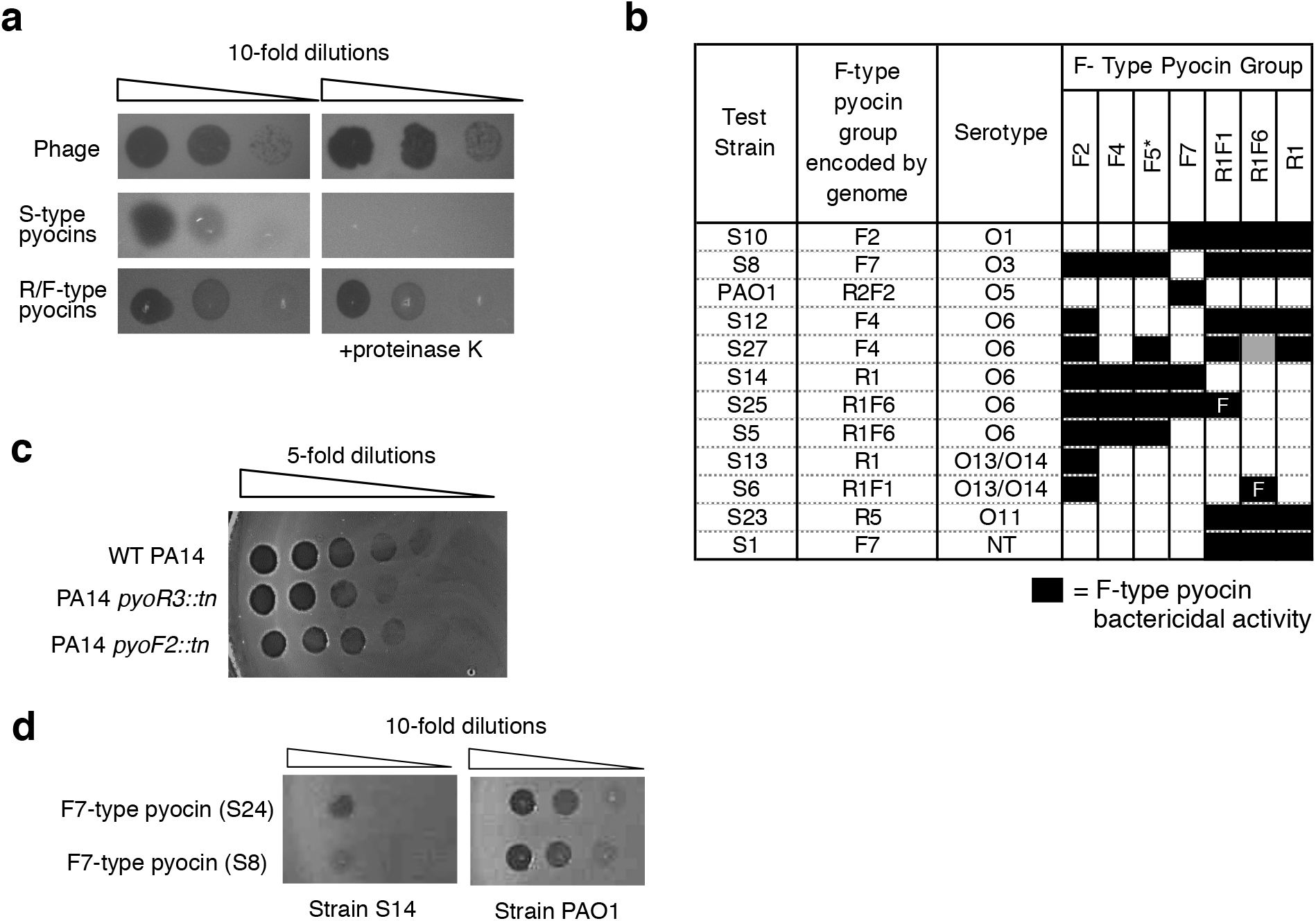
Bactericidal activity of F-type pyocins. (a) Bactericidal activity caused by F- or R-type pyocins can be distinguished from that caused by phages of S-pyocins. S-pyocin activity is destroyed by addition of proteinase K. Zones of clearing resulting from phages resolve into individual plaques upon dilution. (b) The bactericidal activity of F-type pyocins on a selected group of bacterial lawns is shown. These lawns were selected to emphasize the differences in specificity among the different groups. Black boxes denote strains killed by a given F-type pyocin while white boxes denote no killing. The gray box indicates a case where the killing by pyocins was occluded by phage plaquing. Bactericidal activity of the F5 group was determined using a mutant strain of PA14 bearing a transposon insertion in the *pyoR3* gene, so that the lysate contained only F-type pyocin particles (indicated by an asterisk). The F1- and F6-type pyocins were produced in strains that also produced R1-pyocins. By comparing with a strain producing only R1-type pyocins, two strains killed only by these F-type pyocins could be identified (marked with “F”). (c) R- or F-pyocin lysates made from strain PA14 were spotted on a lawn of strain S19. Lysates were produced from wild-type PA14 and strains bearing transposon insertions in either the *pyoR6* or *pyoF10* genes. (d) F7-type pyocins produced from strain S24 or Strain S8 were spotted on lawns of strain S14 or strain PAO1.

All groups of R-type and F-type pyocins identified displayed bactericidal activities against multiple strains. Notably, the killing spectra of lysates were invariably the same if they contained pyocins of the same R- or F-type group (Fig. S5). For example, lysates of four different strains encoding F7 pyocins all displayed bactericidal activity against the same 11 strains (note that in a single case the F-type pyocin activity was occluded by the presence of phage activity as denoted by an orange color, Fig S5). These results demonstrate that our classification of pyocins based on sequence analysis is predictive of biological activity. In Fig. 3b, a small subset of the bactericidal data are shown to emphasize the differences in the killing spectra of the F-type pyocin groups. No two groups kill exactly the same set of bacterial strains; however, considerable overlap exists between some groups like F4 and F5. We also noted that no strain was susceptible to an R- or F-type pyocin that was encoded in its own genome, which is consistent with previous observations that strains are resistant to their own pyocins (35). Since the F1 and F6 groups were encoded only in strains that also encoded R1 pyocins, the killing caused only by the F-type pyocins could not be discerned. However, comparison with results obtained using a strain encoding only an R1 pyocin revealed that strain S25 is susceptible to F1 pyocin as it was killed by a lysate containing F1 and R1 pyocins, but not by a lysate containing only R1 pyocin (Fig. 3b). By the same logic, strain S30 was found to be killed by F6 pyocin. The F5 group was found only in strains that also encode an R-type pyocin. To assess the activity of this group we took advantage of a transposon insertion mutant of an essential R-type pyocin gene in PA14 (36) to detect the activity of the F5 pyocin alone (Fig. 3b).

### F-type and R-type pyocins display similar levels of bactericidal activity

It was previously reported that one R-type pyocin particle is sufficient to kill a single cell, while up to 280 F-type pyocin particles are required to kill the same cell (19). This implies that an F-type pyocin containing lysate would have considerably less killing activity than an R-pyocin containing lysate. However, we observed many cases where lysates of F-type pyocins displayed levels of killing activity as high R-type pyocin lysates. Although R- and F-type pyocin lysates may contain different numbers of particles, we do not expect these numbers to deviate greatly as all pyocin operons utilize the same transcriptional regulatory region. Most convincingly, we tested the bactericidal activity of lysates of two PA14 mutants, one of which carried a transposon insertion in an essential R-type pyocin gene (*pyoR6*) and one of which carried a similar insertion in an essential F-type pyocin gene (*pyoF10*). It can be seen that the bactericidal activity of these two lysates was the same, indicating that F-type pyocins and R-type pyocins are equally lethal to a susceptible host (Fig. 3c). We observed that the same F-type pyocin lysate may display different levels of activity on different strains. For example, lysates of F7 group pyocins displayed greater than 10-fold greater bactericidal activity on strain PAO1 as on strain S14 (Fig. 3d). The previously observed low activity of F-type pyocins was likely caused by use of a non-optimal indicator strain. Overall, our data indicate that F-type pyocins have the potential to kill bacterial cells as efficiently as R-type pyocins.

### The genes downstream of *pyoF10* are required for bactericidal activity

Although homologs of the proteins encoded at the 3’-end of the F-type pyocin cluster (PyoF11 to PyoF15) are encoded in phages and prophages, the roles of these proteins have never been investigated. To determine whether these proteins are essential for bactericidal activity, we tested the activity of F-type pyocin mutants in strain PA14 (group F5). We tested transposon insertion mutations in *pyoF14*, and *pyoF15* from the PA14 non-redundant transposon mutant library (36). We constructed in-frame deletion mutations in *pyoF11* and *pyoF12* and a nonsense mutation in *pyoF13* (Supplementary Materials and Methods). Mutations in each of these genes completely abrogated bactericidal activity, indicating that their protein products play essential roles in the production of functional F-type pyocin particles (Table 2). To ensure that the loss of activity resulting from these mutations was the result of abrogation of only the gene in which the mutation was located, each gene was cloned into a plasmid expression vector (Supplementary Materials and Methods) and we determined whether mutations could be complemented by the plasmid expressed genes. The *pyoF12* mutant could be complemented by a plasmid expressing only *pyoF12* (Table 2). However, complementation of the *pyoF11* mutant required plasmid-based expression of both *pyoF11* and *pyoF12*. A plasmid expressing only *pyoF12* did not complement the *pyoF11* mutant. We conclude that both *pyoF11* and *pyoF12* are essential for bactericidal activity, and that the *pyoF11* in-frame deletion mutation also causes loss of *pyoF12* activity, possibly through a polarity effect. Through a similar series of plasmid based complementation experiments, we determined that *pyoF13, pyoF14*, and *pyoF15* are also essential for bactericidal activity, and that polarity effects are also manifested in this group of genes (Table 2). For example, while the *pyoF15* mutation could be complemented by expression of *pyoF15* alone, complementation of the *pyoF14* mutation required expression of both *pyoF14* and *pyoF15*.

**Table 2.** Plasmid-based complementation of *pyoF11* to *pyoF15* mutants.

| **Mutated Gene** | **Gene(s) expressed on complementing plasmid** | **Complementation Result** |
| --- | --- | --- |
| *pyoF11* | *pyoF11* | No |
| *pyoF11* | *pyoF11+pyoF12* | Yes |
| *pyoF11* | $\Delta pyoF11^1 + pyoF12$ | No |
| *pyoF11* | *pyoF12* | No |
| *pyoF12* | *pyoF12* | Yes |
| *pyoF13* | *pyoF13* | No |
| *pyoF13* | *pyoF13+pyoF14+pyoF15* | Yes |
| *pyoF13* | *pyoF14+ pyoF15* | No |
| *pyoF14* | *pyoF14* | No |
| *pyoF14* | *pyoF14+pyoF15* | Yes |
| *pyoF14* | $\Delta pyoF14^1 + pyoF15$ | No |
| *pyoF15* | *pyoF15* | Yes |
| *pyoF15* | $\Delta pyoF14^1 + pyoF15$ | Yes |

<sup>1</sup>These plasmids express two genes but the first bears an in-frame deletion.

### Serotype correlates with F-type pyocin killing spectra

The outer membrane lipopolysaccharide (LPS) of *P. aeruginosa* is composed of three domains: lipid A, core oligosaccharide and a long-chain polysaccharide O-antigen (37). Most *P. aeruginosa* strains produce two distinct forms of O-antigen; a homopolymer of D-rhamnose known as the common polysaccharide antigen, and a heteropolymer repeat of three to five distinct sugars known as the O-specific antigen (OSA), which forms the basis of *P. aeruginosa* serotyping. Previous studies showed that the OSA acts a receptor for some R-type pyocins, while it blocks killing by other R-type pyocins (26). To investigate the effect of the OSA on the activity of F-type pyocins, we experimentally determined the serotypes of the 30 strains used in this study by a slide agglutination assay. We observed a correlation between the serotype of a strain and its F-type pyocin susceptibility profile (Fig. 4a). For example, all three strains of O2 serotype were resistant to all F-type pyocins, while the four O5 strains were killed only by F7 pyocins. Among the eight O6 serotype strains, the F2 pyocin killed all, but the F4, F5, and F7 pyocins were unable to kill some of these strains (Fig. 4a). The resistance of O6 strains S12 and S27 to the activity of the F4 pyocin is expected as these strains encode an F4 pyocin. However, it is not clear why the F7 pyocin fails to kill O6 strains S12, S27, and S5, or why strain S12, alone among O6 strains, is resistant to F5 pyocin. Similarly, the F1 pyocin kills strain S25 but no other O6 strains, and the F6 pyocin kills strain S6 but no other O13/O14 strains. These data show that factors independent of OSA and pyocin type encoded within a strain contribute to F-type pyocin susceptibility.

**FIG 4.**
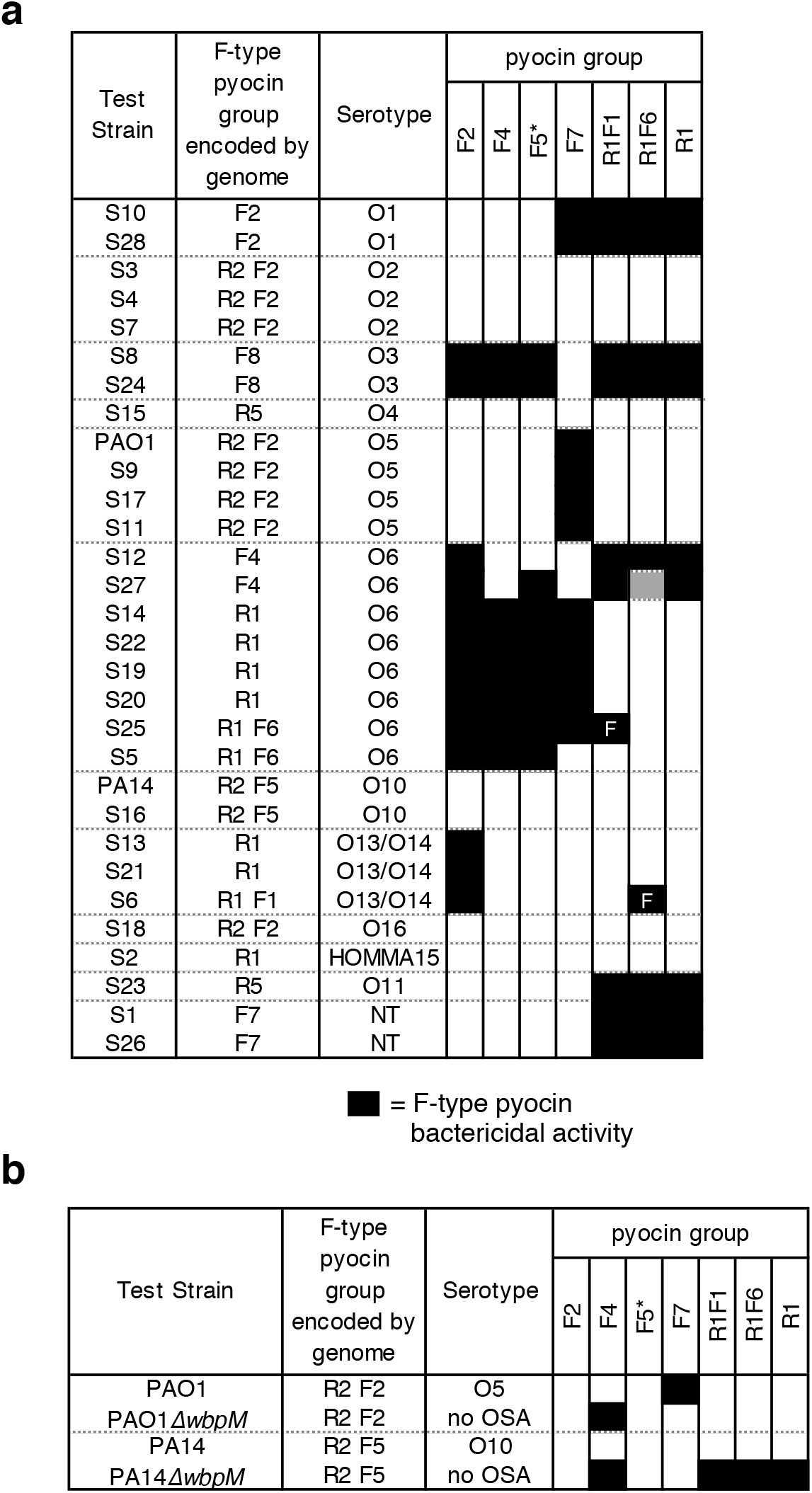
(a) The effect of LPS serotype on bactericidal activity of F-type pyocins. The bactericidal activity of F-type pyocins on a group of bacterial lawns arranged by their serotypes is shown. Black boxes denote strains killed by a given F-type pyocin while white boxes denote no killing. The gray box indicates a case where the killing by pyocins was occluded by phage plaquing. Bactericidal activity of the F5 group was determined using a mutant strain of PA14 bearing a transposon insertion in the *pyoR3* gene, so that the lysate contained only F-type pyocin particles (indicated by an asterisk). The F1- and F6-type pyocins were produced in strains that also produced R1-pyocins. By comparing with a strain producing only R1-type pyocins, two strains killed only by these F-type pyocins could be identified (marked with “F”). (b) The indicated F- type pyocin containing lysates were tested against mutants that lack OSA (*ΔwbpM*).

To directly assess the role of OSA in F-type pyocin activity, we tested Δ*wbpM* mutant strains, which lack OSA in strains PAO1 and PA14 (Fig 4b). The F7-type pyocin is active against PAO1, but was unable to kill the PAO1Δ*wbpM* mutant, suggesting that this pyocin uses the OSA as a receptor. By contrast, the Δ*wbpM* mutants of PAO1 and PA14 became susceptible to the F4 group, though the wild-type strains were not. In this case, the OSA appears to block the pyocin from contacting its receptor. The F2 and F5 groups, which are unable to kill PAO1 or PA14, were also not able to kill the mutants lacking OSA. Strains producing the F1 and F6 group pyocins were unable to kill PAO1 with or without OSA, but PA14Δ*wbpM* did become susceptible to killing. However, this effect may have been due to the R1 pyocins produced by these strains. From these experiments with strains lacking OSA, it is clear that the presence of OSA affects the bactericidal activity of the F4 and F7 groups while the data are inconclusive for the other groups.

### Discovery of new groups of F-type pyocins

To determine if this collection of F-type pyocin described above encompassed the full diversity of F-type pyocins found across the *P. aeruginosa* species, we performed BLAST searches against all *P. aeruginosa* genomes in the NCBI database using a PyoF13 sequence as the query with the goal of identifying homologs with distinct sequences (i.e. share less than 90% sequence identity with those in our established F-type pyocin groups). PyoF13 was chosen for these searches because it is highly conserved among the F-type pyocin operons in its N-terminal region, yet its C-terminal region varies depending on the pyocin group. We discovered three PyoF13 homolog families encoded in F-type pyocin operons that shared less than 70% sequence identity to any other PyoF13 group. F-type pyocins encoding these newly identified PyoF13 varieties were defined as groups F8, F9, and F10. The F9 group is identical to a previously identified group designated as the PA7 group (16). Another identified group, called F11, possessed PyoF13, PyoF14, and PyoF15 homologs that are greater than 95% identical to group F2_2_, F4 and F5, but the PyoF10, PyoF11, and PyoF12 were unique. Finally, group F12 combined PyoF10 to PyoF12 homologs that were 99% identical to group F11 with PyoF13 to PyoF15 homologs that were greater than 95% identical to group F10 (Fig. 5, S1). Group F12 was previously identified in *P. aeruginosa* strain M18 (16). The sequences of proteins PyoF10 to PyoF15 for all eleven F-type pyocin groups can be found in Appendix 1 (Supplementary Material).

**FIG 5.**
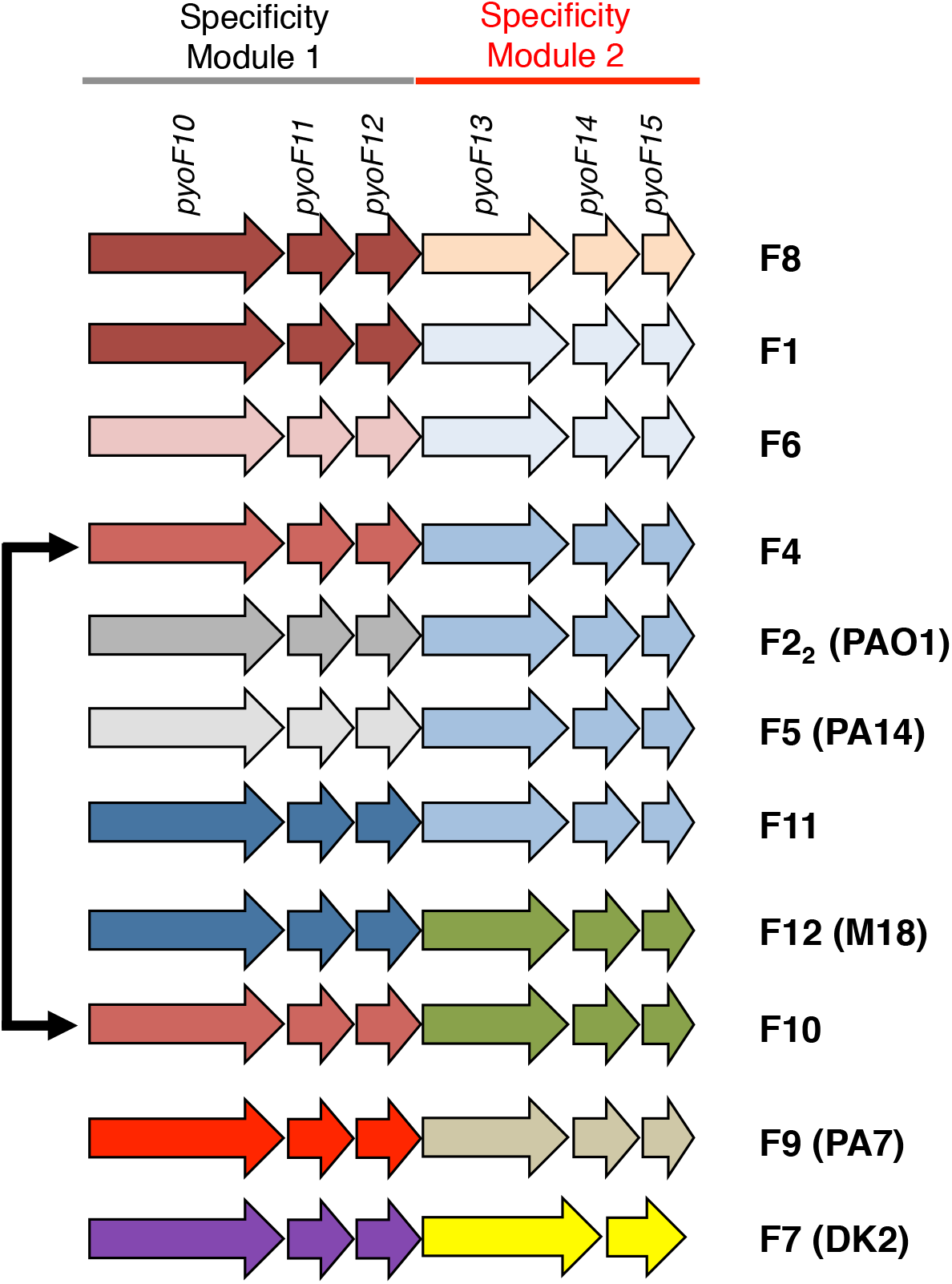
All F-type pyocin groups. A close up of genes encoded at the 3’-end of F-type pyocin clusters shows the 11 different groups identified in our sequenced strains and in the database. Groups of genes are colored the same if the proteins they encode display greater than 90% sequence identity. The extent of Specificity Modules 1 and 2 are shown at the top as are the names of the genes in these regions. *P. aeruginosa* strains where certain groups were previously identified are shown in parentheses.

Pairwise sequence comparisons among all the F-type pyocin groups that we have identified strongly supports the existence of two distinct specificity modules in F-type pyocins (Fig. S1). Whenever the C-terminal regions of PyoF10 proteins in two groups are highly similar (> 90% identity), then the PyoF11 and PyoF12 proteins are also highly similar. Similarly, when two groups have PyoF13 proteins with highly similar C-terminal regions, then the PyoF14 and PyoF15 proteins are also highly similar. Therefore, we have designated regions encoding the C-terminus of PyoF10, PyoF11, and PyoF12 as Specificity Module 1 and regions encoding PyoF13, PyoF14, and PyoF15 as Specificity Module 2. Among our full set of pyocin groups, we observed three instances where the same Specificity Module 1 region assorted with different Specificity Module 2 regions (Fig. 5). We also observed three cases where identical Specificity Module 2 regions assorted with different Specificity Module 1 regions. These data indicate that recombination events have occurred between different F-type pyocin operons.

## DISCUSSION

This study provides a comprehensive analysis of F-type pyocin operons present in *P. aeruginosa* strains. We have defined the conserved genes in these operons and introduced a systematic naming system for them. The initial 21 F-type clusters examined in strains from our collection were categorized into six different groups, two of which were previously known (F1 and F2) and four of which were named in this study (groups F4 to F7). The killing specificity of each of these groups was shown to be distinct. An additional five F-type pyocin groups were discovered bioinformatically, but the killing specificity of these groups remains to be tested. Importantly, we identified two highly diverse F-type pyocin genes, *pyoF11* and *pyoF12*, which are not annotated as genes in many *P. aeruginosa* strains, yet are essential for bactericidal activity. The sequence diversity in these two genes contributes to defining the F-type pyocin groups. An important finding is that there are no genes in the F-type pyocin operons that are not also found in the genomes of phages or prophages. Thus, the ability of these pyocins to efficiently kill bacteria while isolated phage tails do not must be due to sequence modifications within their phage-derived proteins, not to the presence of unique toxin-encoding genes (unless such genes are encoded elsewhere in the *P. aeruginosa* genome). Fully active R-type pyocins have been produced heterologously in *E. coli* from a plasmid vector including only genes from the R-type pyocin operon, indicating that their toxicity does not rely on genes outside of this region (9).

Analysis of the F-type pyocin operons clearly indicates the genes that are involved in killing specificity. The proteins encoded by the *pyoF2* to *pyoF9* genes are highly similar in all the F-type pyocin groups. Divergence among the groups begins with the last 60 residues of PyoF10 and extends through PyoF15. We defined the F-type pyocin groups according to sequence identity among these proteins (Fig. 5). Since F-type pyocins within the same group invariably displayed the same killing spectra (Fig. S5), we conclude that some or all of the *pyoF10* to *pyoF15* region determines killing specificity. By examining the patterns of recombination among the specificity genes, we defined two specificity modules: Module 1 (*pyoF10* to *pyoF12*) and Module 2 (*pyoF13* to *pyoF15*). The occurrence of highly similar Module 1 regions with distinct Module 2 regions and vice versa in different F-type pyocin groups indicates that the two modules act independently of one another (Fig. 5). This conclusion is supported by the appearance of *pyoF11* and *pyoF12* homologs without adjacent *pyoF13*-*pyoF15* homologs and vice versa in phages and prophages. Since these families of proteins have not been characterized, defining their roles in host specificity will be an important goal for further study.

The strong sequence similarity between most of the F-type pyocin genes implies that all the groups are descended from a common ancestor that likely arose from a defective prophage. However, it is also clear that some type of horizontal gene transfer mechanism has been responsible for the evolution of the specificity regions, which are comprised of different combinations of Specificity Modules 1 and 2. These reassortments could be caused by phages carrying genes that are similar to these F-type pyocin genes occasionally recombining with the homologous F-type pyocin genes. With respect to the evolution of the F- and R-type pyocins as a whole, it is relevant that the F-type display considerably more divergence among their specificity-determining genes as compared to the R-type (Fig. S1, S4). This suggests that the F-type pyocins may have arisen first and, thus, have had more time to diverge. Also supporting the possibility that the F-type pyocin operon appeared first is that the R-type pyocin genes are inserted in the middle of the lysis genes, which comprise an intact lysis cassette in strains possessing only an F-type operon.

The bacterial cell surface receptors of F-type pyocins were previously unknown. Our examination of the activity of the different F-type pyocin groups on 12 different serotypes of *P. aeruginosa* revealed a clear correlation between bactericidal activity and O-antigen serotypes (Fig. 4a). Further supporting a role for the OSA in the activity of at least some F-type pyocins is the fact that activity of the F7 group required the presence of OSA, while group F4 activity was blocked by OSA (Fig. 5b). While the OSA serotype clearly influences F-type pyocin host recognition, this is not the only determining factor. For example, the F7 pyocin is able to kill some but not all strains with the O6 serotype. In addition, F-type pyocins of a given type were consistently unable to kill strains encoding the same type of F-type pyocin, regardless of serotype. The mechanism of this self-immunity is not known. Many bacteriocins, such as S-type pyocins and colicins, are encoded with specific immunity proteins (11). However, there is no obvious immunity protein candidate encoded within F-type pyocin clusters as each gene is homologous to phage tail proteins. It is possible that no specific immunity proteins exist for R-type or F-type pyocins. Rather, strains may have evolved to resist their resident R- and F-type pyocins by altering their cell surface in subtle ways undetectable by the antibodies used in serotyping.

Overall, our study shows that F-type pyocins are produced by a large number of *P. aeruginosa* strains, they all possess antimicrobial properties, and they are promising candidates to study for the development of new therapeutics. Our identification of the specificity determinants of F-type pyocins points the way toward precisely engineering their killing as has been done with the contractile R-type pyocins and non-contractile tailocins of *Listeria (7, 8, 10)*.

## MATERIALS AND METHODS

### Whole genome sequencing

Genomic DNA was isolated using a genomic DNA extraction kit (Bio Basic Inc). Next-generation whole genome sequencing was performed by the Donnelly Sequencing Center, University of Toronto, using Illumina HiSeq2500. *De novo* assembly of reads into contigs was performed using Velvet version 2.2.5 (38). Genes *trpE* and *trpG* were located and the region between these genes was analyzed using Geneious (39).

### Bioinformatic analysis

Most of the bioinformatic analyses, including BLAST (40) searches and genome synteny analyses were carried out on a custom database comprised of 755 tailed phage genomes and 2,119 bacterial genomes downloaded from the National Center for Biotechnology Information (NCBI) Refseq database in April 2013. This database contains diverse phage and bacterial species, but was small enough to allow manual analysis of the protein sequences and the genomic context of genes encoding proteins related to pyocin proteins. This work was aided by a synteny viewing and phage gene annotation toolkit developed in our laboratory, which will be described in detail elsewhere. Sequence alignment analysis was performed in Jalview (41). To identify protein sequences similar to less frequently occurring proteins found in the pyocin cluster (e.g. PyoR1, PyoF11, and PyoF12), alignments were constructed of the pyocin proteins. HMMER3 (32) was then used to create Hidden Markov Models (HMMs). These HMMs were used to detect proteins similar to a given pyocin protein. The genome context of genes encoding these similar proteins within phage genomes was assessed to support a conclusion that the pyocin protein possesses the same function as the phage protein. BLAST searches to identify new groups of F-type pyocins were carried out against all *P. aeruginosa* genomes available at NCBI in April, 2018.

HHpred searches were carried out using the online server (https://toolkit.tuebingen.mpg.de/hhpred) (42). HMM-based searches were carried out using HMMer (32) and analyzed by searching the Pfam (43) and TIGRfam (www.jcvi.org/cgi-bin/tigrfams/index.cgi) databases.

### Transmission electron microscopy

A continuous carbon film coated EM grid was made hydrophilic by glow discharge. 5 μl of sample was applied to the surface of the grid and left for absorption for 2 minutes. Excess sample was blotted away using the corner of a filter paper. The grid was washed three times with water and stained with 2% (w/v) uranyl acetate. Grids were examined with a Hitachi H-7000 microscope.

### Assays of pyocin and phage bactericidal activity

To generate lysates containing pyocins and/or phages, 5 ml cultures started from overnights were grown in LB at 30 °C until the cells reached an OD_600_ of 0.4. Mitomycin C, was then added to a final concentration of 2 μg/ml and shaking at 30 °C was resumed for 3 h or until cell lysis occurred. Chloroform was added to all induced cultures (1-2 drops/ml) to ensure maximum bacterial lysis. In experiments testing complementation from plasmids, 0.2% arabinose was added to cells after 1 h of growth at 30 ^°^C to induce the expression of proteins from the plasmid prior to addition to mitomycin C to induce F-type pyocin induction from the genome. After lysis, cultures were incubated at room temperature with DNase (10 μg/ml) for 30 min prior to centrifugation at 10000 rpm for 10 min. For activity assays, 2 µl volumes of dilutions of these lysates were spotted onto lawns of *P. aeruginosa* strains. Lawns of strains to be tested were made by adding 150 μl of overnight culture to 3 ml of molten 0.7% top agar, which was immediately poured onto an LB agar plate and allowed to harden. To distinguish S-type pyocin activity, duplicate lawns were poured containing proteinase K (100 μg/ml). At least three biological replicates were performed for each strain and lysate combination. A lysate was scored as positive if it displayed a strong zone of cell growth inhibition in every assay. There was a range of activity in positive lysates with some displaying moderate zones of growth inhibition even when diluted 10^2^-fold and others displaying strong activity only when undiluted. Lysates scored as negative displayed very weak or no zones of growth inhibition in all replicate assays.

### Serotyping of *P. aeruginosa* strains

Strains were serotyped using the slide agglutination method using commercial antisera (MAST Diagnostics) against all 20 *P. aeruginosa* serotypes recognized by the International Antigenic Typing Scheme (37).

## ACKNOWLEDGEMENTS

This work was supported by operating grants from the Canadian Institutes of Health Research (CIHR) to A.R.D. (XNE-86943 and FDN-15427), K.L.M (MOP-136845), and A.W.E. (PHT-148819). S.S. was supported by a CIHR Doctoral Scholarship.

## SUPPLEMENTAL MATERIAL

## Supplemental Materials and Methods

In separate file.

## Supplementary Table 1

List of strains used and origin (in separate file)

## Supplementary Table 2

List of primers used (in separate file)

## Supplementary Figure Legends

**FIG S1** Pairwise identities among F-type pyocin proteins encoded at the 3’-end of the operon. All against all pairwise percent identities for the indicated proteins are shown. The F-pyocin groups of the proteins that are being compared are indicated at the sides and top of each table. Numbers that are shaded denote groups that share Specificity Modules. The PyoF10 comparisons include only the C-terminal 60 amino acids. PyoF13 comparisons include only the last 210 residues of these proteins. The F7 group does not encode PyoF14, so these boxes are left blank. “<20” denotes sequences that could not be well aligned in a pairwise alignment. F21 and F22 refer to the duplicated PyoF13, PyoF14, and PyoF15 proteins encoded in the F2 group.

**FIG S2** Protein sequence alignments of PyoF11 and PyoF12 from each F-type pyocin group. (a) An alignment of PyoF11 homologs is shown from the 11 F-type pyocin groups and selected phages. (b) An alignment of PyoF11 homologs is shown from the 11 F-type pyocin groups and selected phages. The phage proteins are from *Burkholderia* phages KS9 (NC_013055) and BcepGomr (NC_009447); *P. aeruginosa* phages LIT1 (NC_013692) and LUZ7 (NC_013691); and *E. coli* phages T1 (NC_005833) and N15 (NC_001901).

**FIG S3** Protein sequence alignments of PyoF13 from each F-type pyocin group. (a) An alignment of the N-terminal 140 amino acids of PyoF13 homologs from the 11 F-type pyocin groups is shown. (b) An alignment of the C-terminal 210 amino acids of PyoF13 homologs from the 11 F-type pyocin groups is shown.

**FIG S4** Protein sequence alignments of the tail fiber proteins (PyoR6) from each R-type pyocin group. (a) An alignment of the N-terminal 450 amino acids of the PyoR6 homologs is shown. (b) An alignment of the C-terminal 250 amino acids PyoR6 homologs is shown. (c) The pairwise sequence identities of the PyoR6 N-terminal and C-terminal regions are shown. The locus tags for the proteins shown are R1 (PLES_06171), R2 (PA14_08050), and R5 (PA0620).

**FIG S5** The bactericidal activity of all F-type pyocin lysates tested on all of the *P. aeruginosa* strains used in this study.

